# Changes in rumen microbiota of cows in response to dietary supplementation with nitrate, linseed and saponin alone or in combination

**DOI:** 10.1101/383067

**Authors:** Milka Popova, Jessie Guyader, Mathieu Silberberg, Ahmad Reza Seradj, Cristina Saro, Aurélien Bernard, Christine Gérard, Cécile Martin, Diego P Morgavi

**Affiliations:** Université Clermont Auvergne, INRA, VetAgro Sup, UMR Herbivores, F-63122 Saint-Genès-Champanelle, France; NEOVIA, Talhouёt, F-56250 Saint Nolff, France; Departament Producció Animal, Universidad de Lleida, ETSEA, Alcalde Rovira Roure 191, 25198, Lleida, Spain

**Author notes:** Address correspondence to Milka Popova.

## Abstract

Dietary supplementation with linseed, saponins and nitrate is a promising methane mitigation strategy in ruminant production. The main objective of this work was to assess the effects of these additives on the rumen microbiota in order to understand underlying microbial mechanisms of methane abatement. Two 2 × 2 factorial design studies were conducted simultaneously, which also allowed us to make a broad-based assessment of microbial responses. Eight non-lactating cows were fed diets supplemented with linseed or saponin in order to decrease hydrogen production and nitrate to deviate hydrogen consumption; also, combinations of linseed plus nitrate or saponin plus nitrate were used to explore the interaction between dietary treatments. Amplicon sequencing of 18S and 16S rRNA genes was employed to characterise rumen microbes. Nitrate fed alone or in combination in both studies dramatically affected the composition and structure of rumen microbiota, though impacts were more evident in one of the studies. Linseed moderately modified bacterial community structure with no effect on rumen methanogens and protozoa. Indicator OTU analysis revealed that both linseed and nitrate reduced the relative abundance of hydrogen-producing *Ruminococcaceae*. Linseed increased the proportion of bacteria known to reduce succinate to propionate, whereas nitrate supplementation increased nitrate-reducing bacteria and decreased the metabolic activity of rumen methanogens. Saponins had no effect on the microbiota. Inconsistency found between the two studies, when nitrate was fed to the cows could be explained by changes in microbial ecosystem functioning rather than changes in microbial community structure.

**Importance:** This study aimed at identifying the microbial mechanisms of enteric methane mitigation when linseed, nitrate and saponins were fed to non-lactating cows alone or in a combination. Hydrogen is a limiting factor in rumen methanogenesis. We hypothesised that linseed and saponins would affect hydrogen producers and nitrate would deviate hydrogen consumption thus leading to reduced methane production in the rumen. Contrary to what was foreseen, both linseed and nitrate had a deleterious effect on hydrogen producers; linseed also redirected hydrogen consumption towards propionate production, whereas nitrate stimulated the growth of nitrate reducing and hence hydrogen-consuming bacterial taxa. Fundamental knowledge of microbial mechanism involved in rumen methanogenesis, provides novel insights for the development of new or the optimisation of existing methane mitigation strategies.

## Introduction

Methane emissions associated with ruminant livestock production are an important contributor to global greenhouse gas emissions (1). Rumen methanogenesis is a naturally occurring process that involves methanogenic archaea consuming hydrogen to reduce carbon dioxide. Hydrogen and carbon dioxide production occurs during feed fermentation by bacteria and protozoa; hydrogen availability is a limiting factor for methane production. In addition, there is a significant linear relationship between protozoa concentration in the rumen and methane emissions (2). Among the measures that have been undertaken to reduce methane production by ruminants, diet composition and inclusion of feed additives have received the most attention (3). Among them, nitrate added to ruminants’ diets consistently and persistently lowers methane emissions (4). Linseed oil, which is rich in linoleic acid, has proven to be one of the most efficient lipid sources used in methane mitigation strategies (4). Saponins are natural phytogenic feed additives used to improve animal feeding and production characteristics (5). Theoretically, these three additives lead to decreased methane production via different modes of action. Nitrate is an alternative electron acceptor as its reduction competes with methane production for hydrogen (6). Additionally, nitrate or its reduced forms might be toxic to rumen methanogens and protozoa (7), but this effect was not systematically reported (8, 9). Lipids from linseed (and fats in general) added to diets replace a proportion of dietary carbohydrates and, as rumen microbes do not ferment them, less hydrogen is produced. Protozoal numbers have been reported to decrease with supplementary linseed oil (8, 10), though this effect was not always observed (11). Saponins can reduce methanogenesis by a toxic effect on rumen protozoa (5), but *in vivo* results are contrasting as rumen microbes can deglycosylate and thus inactivate saponins (12).

Based on available information, we hypothesised that linseed oil and saponins would mainly affect hydrogen production (by a toxic effect on protozoa or by providing alternative substrate for rumen fermentation) and nitrate would mainly modulate hydrogen consumption pathways (by providing an alternative hydrogen sink). We performed amplicon-type sequencing analysis of rumen contents, sampled during two previous studies (8, 13); the first one reported the effect of linseed, nitrate and linseed plus nitrate supplementation on enteric methane production; tea saponin replaced linseed in the second one. The primary aim of the current study was to search for changes in rumen microbiota structure and methanogenic activity that could explain observed reductions in methane emissions.

Minor but significant changes induced by treatment can be masked by spurious between-group differences related not to the treatment, but rather to the host animal, the diet, or sample management. Moreover, it is not unusual to find reports on nitrate and fatty acid supplementation where methane decreased in a similar way, but effects on rumen microbiota were contrasting (14-17). On the other hand, it was recently shown that combination of microbial data from multiple sets of microbially related organisms should increase specificity and allow identification of causal microbes (18). Therefore, we took advantage of the data available from two independent studies, analysed it separately but following the same procedures and made an integrated interpretation. Our secondary objective was to try to find clues to explain inconsistency in results from published studies.

## Results

Eight non-lactating dairy cows were randomly allocated to two 2 × 2 factorial designs. In Study 1, dietary treatments consisted of control (CTL) diet, supplemented alternatively with linseed oil (LIN), nitrate (NIT) and linseed plus nitrate (LIN+NIT); in Study 2, tea saponin (TEA) replaced linseed oil. In order to achieve adequate statistical power, the statistical model for both studies included cow as random effect and fixed effects were experimental period and i) in Study 1 linseed (CTL and NIT *versus* LIN and LIN+NIT), nitrate (CTL and LIN *versus* NIT and LIN+NIT) and their interaction linseed*nitrate and ii) in Study 2, saponin (CTL and NIT *versus* TEA and TEA+NIT), nitrate (CTL and TEA *versus* NIT and TEA+NIT in Study 2) and their interaction saponin*nitrate. Throughout the text linseed, nitrate and saponin will refer to diet contrasts detailed above.

In Study 1, compared to CTL, dietary treatments LIN, NIT and LIN+NIT decreased methane production (g/day) by 22%, 29% and 33% respectively and methane yield (g/kg dry matter intake) by 25%, 29% and 32% (8). In Study 2, NIT and TEA+NIT decreased methane production by 42% and 34% and methane yield by 36% and 29% respectively, compared to CTL (13). TEA alone had no effect on methane production or on volatile fatty acid (VFA) profiles.

In both studies, *Bacteroidales* and *Clostridiales* were the dominant bacterial orders and accounted for more than 88% of the classified reads, regardless of the dietary treatment (Figure S1). Sequences affiliated with the *Methanobrevibacter* genus accounted for 80% of all archaea sequences in both studies, followed by *Methanosphaera*, unclassified methanogens and three *Methanomassiliicoccaceae* genera (Figure S1).

### Linseed moderately affected bacterial community composition with no effect on rumen methanogens and protozoa

Non-metric multidimensional scaling did not reveal any distinct clustering of bacterial communities (Figure 1) and total bacterial numbers were similar (Table 1) in cows receiving or not linseed-supplemented diets. However, the richness index was reduced by the linseed treatment (Table S1) and linseed increased (P<0.05) relative abundance of *Selenomonadales, Synergistales, Elusimicrobiales* and *Micrococcales* (Table 2). Moreover, indicator species analysis showed that *Ruminococcaceae*−related OTUs characterised the bacterial community of cows not receiving linseed supplementation (Figure 2, Table S2).

**Figure 1.**
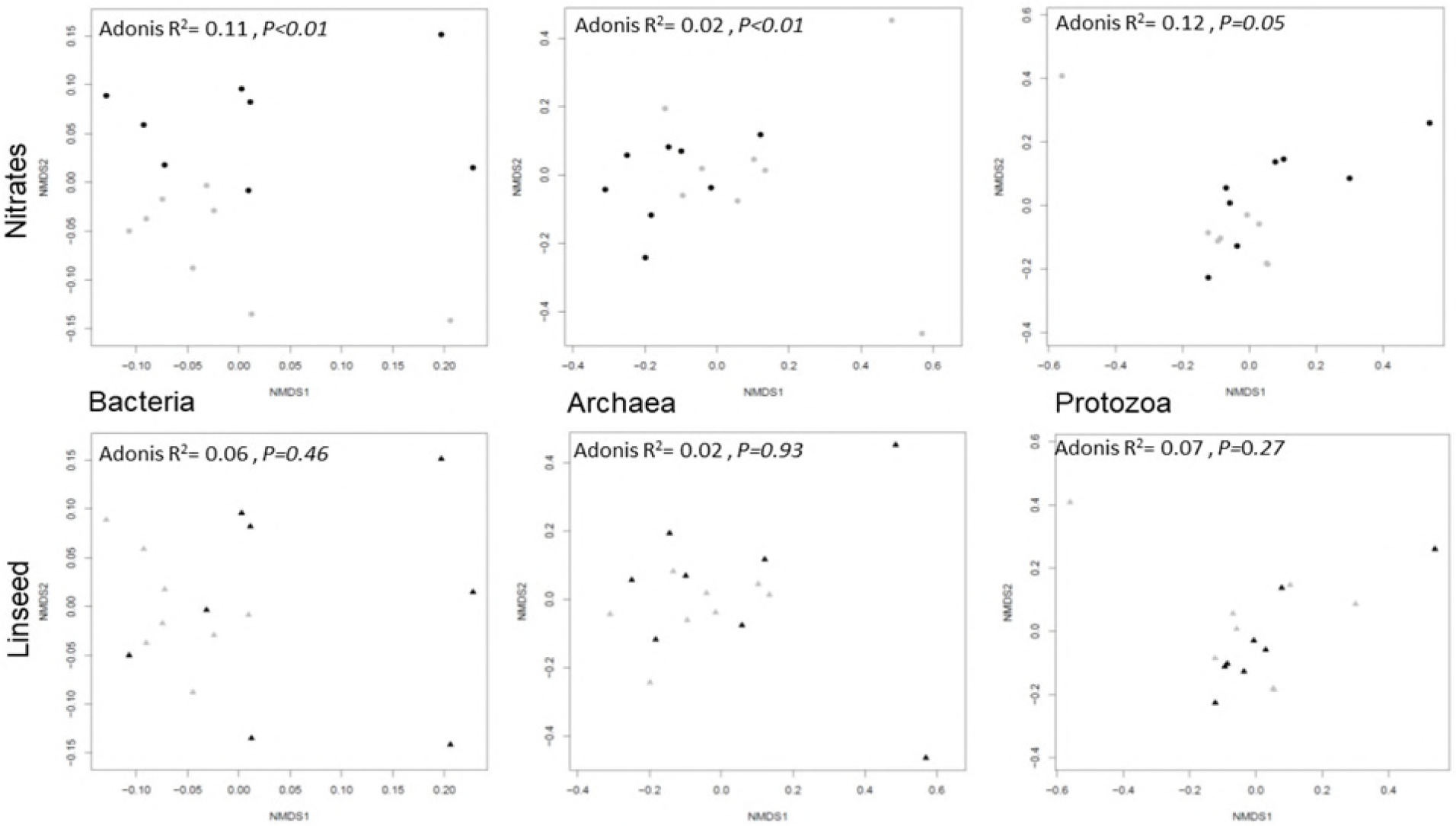
Structure and composition of bacterial, archaeal and protozoal communities in Study 1, related to nitrate or linseed treatments (black symbols) and respective controls (grey symbols), were examined by multivariate analysis. Non-metric multidimensional scaling plots derived from Bray Curtis dissimilarities between cows. Each symbol is representative of a single cow. Samples are plotted along the first two component axes. Microbial composition was compared using Adonis.

**Figure 2.**
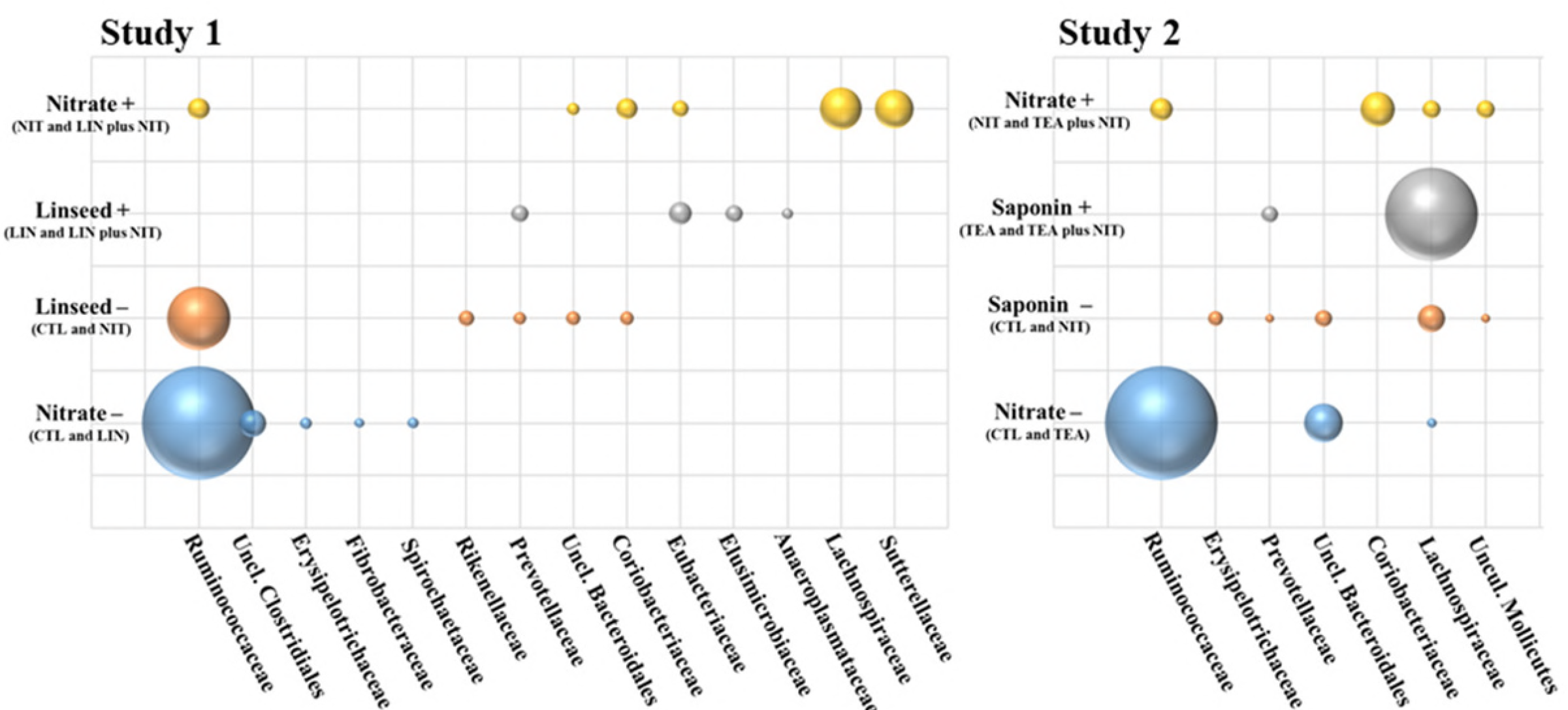
Bubble charts showing indicator OTU distribution by dietary treatment in the rumen of non-lactating cows fed methane-reducing additives. Bubble size reflects the count number in the rarefied dataset.

**Table 1.**
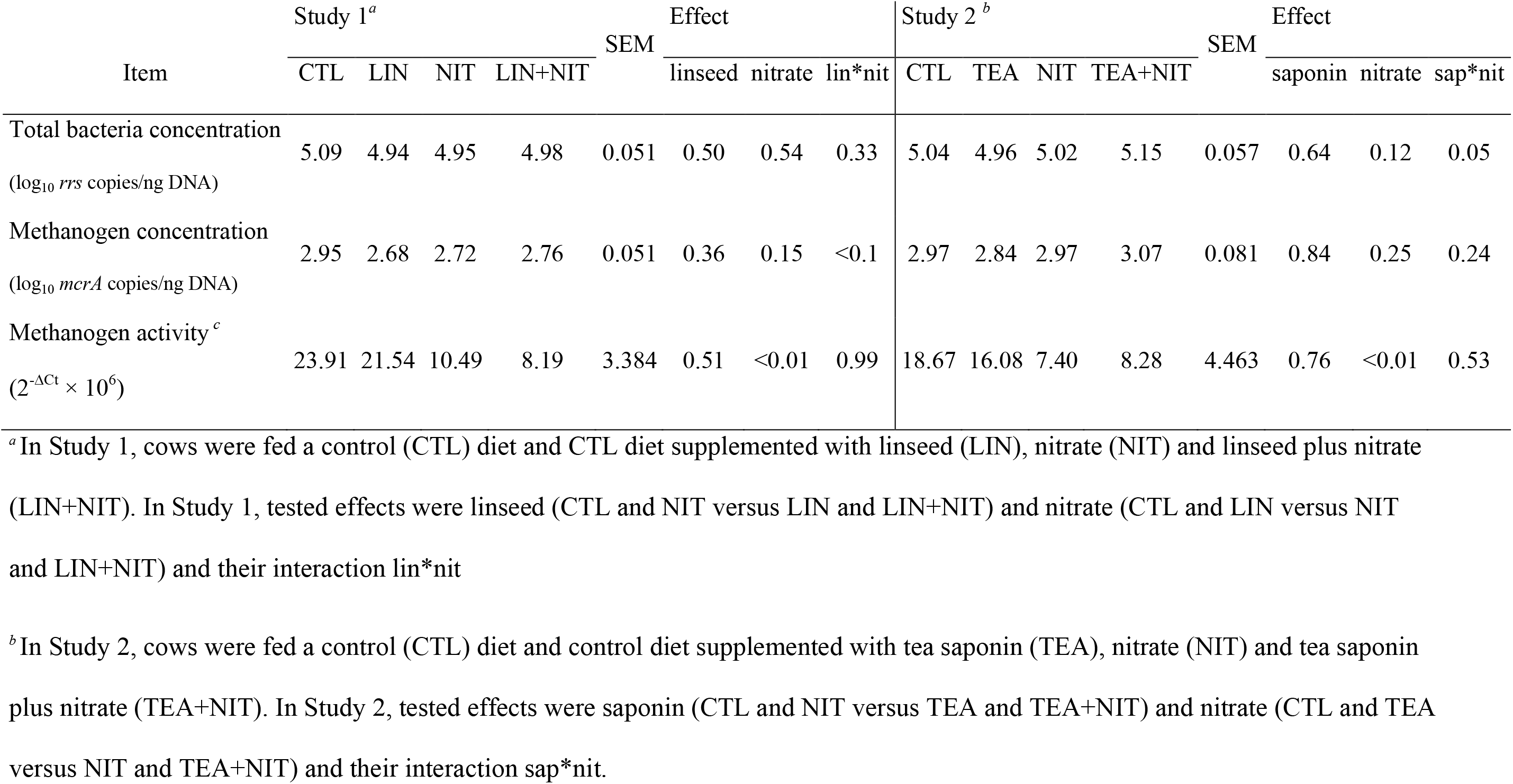
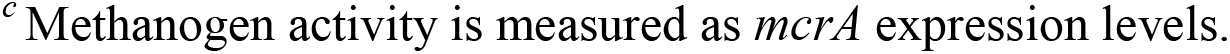
Abundance of total bacteria, and abundance and activity of methanogenic archaea in the rumen of non-lactating cows fed methane-reducing additives.

| Item | Study 1<sup>a</sup> |  |  |  | SEM | Effect |  |  |  | Study 2<sup>b</sup> |  |  |  | SEM | Effect |  |  |
| --- | --- | --- | --- | --- | --- | --- | --- | --- | --- | --- | --- | --- | --- | --- | --- | --- | --- |
|  | CTL | LIN | NIT | LIN+NIT |  | linseed | nitrate | lin*nit |  | CTL | TEA | NIT | TEA+NIT |  | saponin | nitrate | sap*nit |
| Total bacteria concentration<br>(log<sub>10</sub> *rrs* copies/ng DNA) | 5.09 | 4.94 | 4.95 | 4.98 | 0.051 | 0.50 | 0.54 | 0.33 |  | 5.04 | 4.96 | 5.02 | 5.15 | 0.057 | 0.64 | 0.12 | 0.05 |
| Methanogen concentration<br>(log<sub>10</sub> *mcrA* copies/ng DNA) | 2.95 | 2.68 | 2.72 | 2.76 | 0.051 | 0.36 | 0.15 | <0.1 |  | 2.97 | 2.84 | 2.97 | 3.07 | 0.081 | 0.84 | 0.25 | 0.24 |
| Methanogen activity<sup>c</sup><br>(2<sup>-ΔCt</sup> × 10<sup>6</sup>) | 23.91 | 21.54 | 10.49 | 8.19 | 3.384 | 0.51 | <0.01 | 0.99 |  | 18.67 | 16.08 | 7.40 | 8.28 | 4.463 | 0.76 | <0.01 | 0.53 |

<sup>a</sup>In Study 1, cows were fed a control (CTL) diet and CTL diet supplemented with linseed (LIN), nitrate (NIT) and linseed plus nitrate (LIN+NIT). In Study 1, tested effects were linseed (CTL and NIT versus LIN and LIN+NIT) and nitrate (CTL and LIN versus NIT and LIN+NIT) and their interaction lin\*nit
<sup>b</sup>In Study 2, cows were fed a control (CTL) diet and control diet supplemented with tea saponin (TEA), nitrate (NIT) and tea saponin plus nitrate (TEA+NIT). In Study 2, tested effects were saponin (CTL and NIT versus TEA and TEA+NIT) and nitrate (CTL and TEA versus NIT and TEA+NIT) and their interaction sap\*nit.
<sup>c</sup> Methanogen activity is measured as *mcrA* expression levels.

**Table 2.**
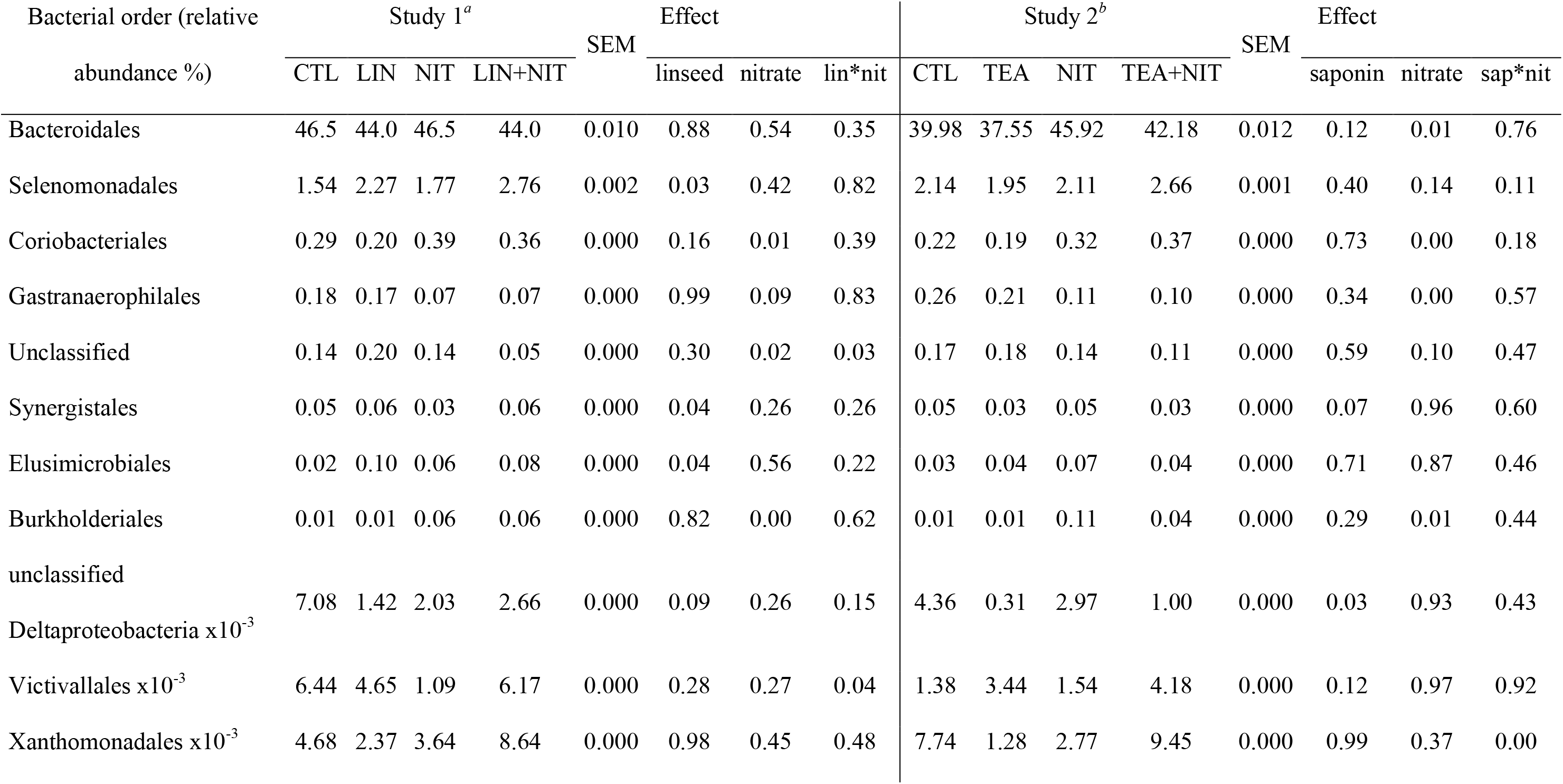
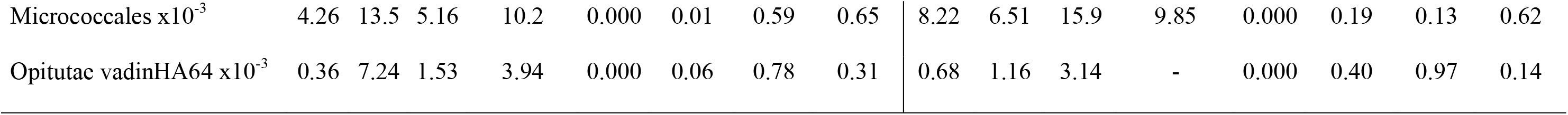
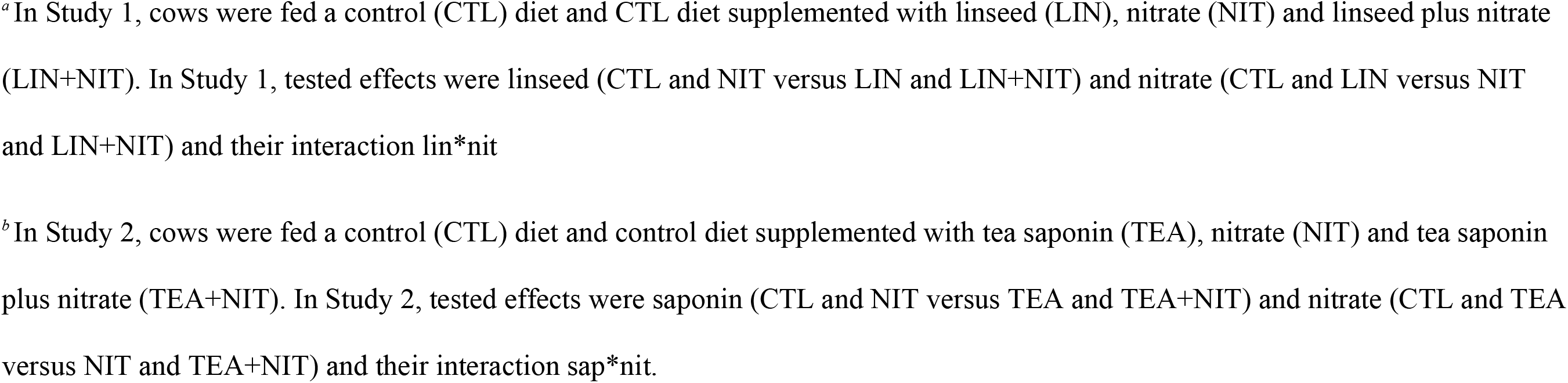
Bacterial orders significantly affected by at least one dietary treatment in the rumen of non-lactating cows fed methane-reducing additives. Values are the mean of four observations and analysis was performed on square-transformed taxonomic tables using the aov function in R.

Regarding methanogen concentration, *mcrA* copy numbers per ng extracted DNA were not affected by linseed supplementation (Table 1), and neither was overall community structure (Figure 1, Table 3).

**Table 3.** Archaeal genera detected in the rumen of non-lactating cows fed methane-reducing additives. Values are the mean of four observations and analysis was performed on square-transformed taxonomic tables using the aov function in R.

| Archaeal genus<br>(relative abundance %) | Study 1<sup>a</sup> |  |  |  | SEM | Effect |  |  | Study 2<sup>b</sup> |  |  |  | SEM | Effect |  |  |
| --- | --- | --- | --- | --- | --- | --- | --- | --- | --- | --- | --- | --- | --- | --- | --- | --- |
|  | CTL | LIN | NIT | LIN+NIT |  | linseed | nitrate | lin*nit | CTL | TEA | NIT | TEA+NIT |  | saponin | nitrate | sap*nit |
| Methanobrevibacter | 80.12 | 82.13 | 80.05 | 84.64 | 0.008 | 0.31 | 0.72 | 0.70 | 78.96 | 79.25 | 79.41 | 78.92 | 0.001 | 0.96 | 0.97 | 0.86 |
| Methanosphaera | 9.10 | 6.83 | 9.78 | 8.97 | 0.005 | 0.62 | 0.75 | 0.87 | 9.98 | 9.28 | 6.61 | 12.24 | 0.008 | 0.15 | 0.75 | 0.10 |
| Unclassified methanogens | 4.59 | 4.66 | 3.37 | 2.52 | 0.004 | 0.40 | 0.01 | 0.35 | 5.18 | 5.15 | 5.92 | 4.11 | 0.003 | 0.45 | 0.82 | 0.37 |
| Uncl. Methanomassiliicoccaceae | 2.96 | 3.12 | 3.87 | 1.58 | 0.003 | 0.11 | 0.61 | 0.13 | 2.74 | 1.51 | 3.44 | 1.67 | 0.003 | 0.06 | 0.63 | 0.94 |
| Group12 | 1.99 | 2.21 | 2.50 | 2.05 | 0.001 | 0.75 | 0.99 | 0.65 | 1.51 | 2.41 | 3.27 | 2.36 | 0.003 | 0.76 | 0.19 | 0.15 |
| Group10 | 0.60 | - | 0.16 | - | 0.001 | 0.08 | 0.52 | 0.52 | 0.99 | 0.84 | 0.61 | 0.56 | 0.001 | 0.86 | 0.46 | 0.75 |
| Methanomicrobium | 0.24 | 0.08 | 0.01 | - | 0.000 | 0.34 | 0.11 | 0.56 | 0.51 | 0.46 | 0.62 | 0.04 | 0.001 | 0.14 | 0.64 | 0.45 |
| Group8 | 0.10 | 0.09 | 0.03 | - | 0.000 | 0.70 | 0.27 | 0.55 | 0.02 | 0.00 | 0.00 | - | 0.000 | 0.12 | 0.04 | 0.37 |
| Other Euryarchaeota | 0.09 | 0.07 | 0.00 | 0.01 | 0.000 | 0.93 | 0.13 | 0.90 | 0.00 | 0.23 | - | 0.01 | 0.000 | 0.07 | 0.15 | 0.20 |
| Other Methanobacteriaceae | 0.07 | 0.05 | 0.08 | 0.05 | 0.000 | 0.42 | 1.00 | 0.81 | 0.04 | 0.05 | 0.04 | 0.06 | 0.000 | 0.43 | 0.90 | 0.74 |
| Methanobacterium | 0.04 | 0.71 | 0.03 | 0.01 | 0.001 | 0.25 | 0.09 | 0.16 | 0.01 | 0.56 | 0.05 | - | 0.001 | 0.24 | 0.11 | 0.07 |
| Group9 | 0.03 | 0.02 | 0.09 | 0.16 | 0.000 | 0.97 | 0.33 | 0.82 | 0.04 | 0.21 | 0.03 | 0.02 | 0.000 | 0.05 | 0.00 | 0.02 |
| Group11 | 0.02 | 0.01 | 0.02 | 0.01 | 0.000 | 0.19 | 0.84 | 0.97 | 0.01 | 0.02 | 0.01 | 0.01 | 0.000 | 0.25 | 0.91 | 0.19 |
| Other Methanococcales | 0.01 | - | - | - | 0.000 | 0.31 | 0.31 | 0.31 | - | 0.01 | - | - | 0.000 | 0.31 | 0.31 | 0.31 |
| Other Archaea | 0.00 | - | - | - | 0.000 | 0.36 | 0.36 | 0.36 | 0.02 | - | - | 0.00 | 0.000 | 0.43 | 0.43 | 0.23 |

Feeding linseed did not modify protozoa community structure and composition, as compared to the respective control treatment (Figure 1, Table 4). There were 3 indicator OTUs identified, 2 associated with CTL diet and one with LIN diet, but they all represented less than 0.01% of the rarefied dataset (13 809 reads per individual)

**Table 4.**
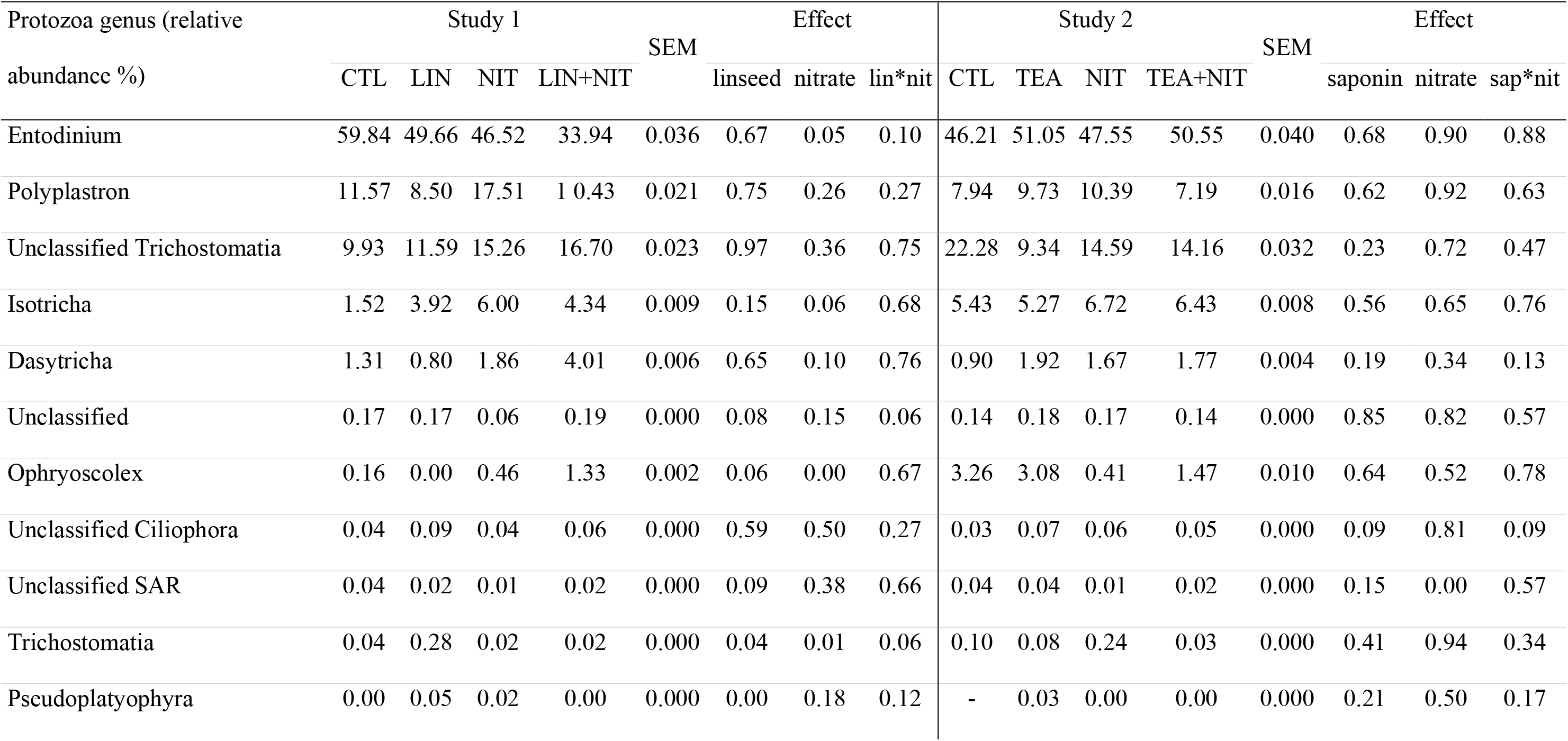
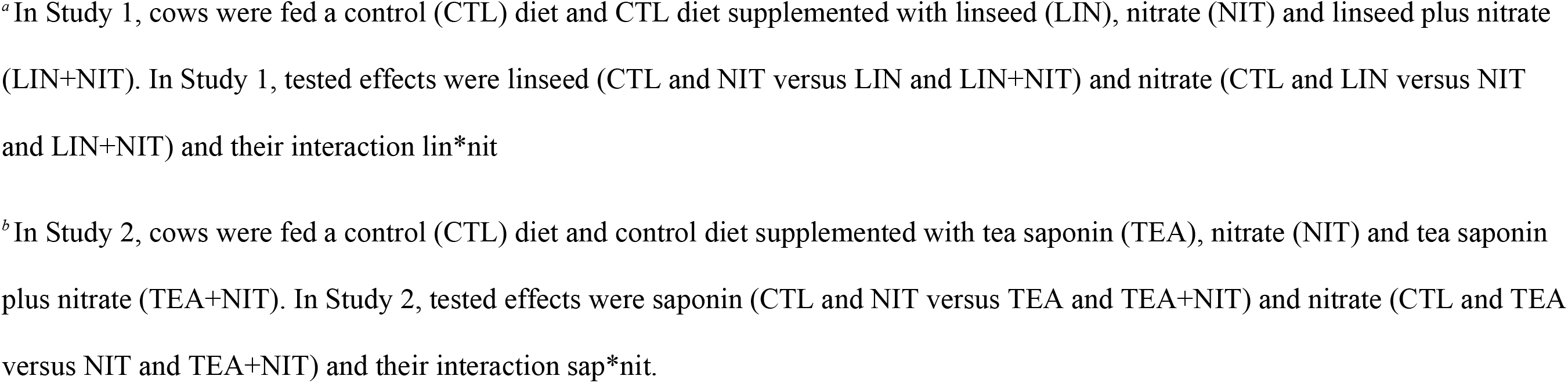
Protozoa genera detected in the rumen of non-lactating cows fed methane-reducing additives. Values are the mean of four observations and analysis was performed on square-transformed taxonomic tables using the aov function in R.

| Protozoa genus (relative<br>abundance %) | Study 1 |  |  |  | SEM | Effect |  |  |  | Study 2 |  |  |  | SEM | Effect |  |  |
| --- | --- | --- | --- | --- | --- | --- | --- | --- | --- | --- | --- | --- | --- | --- | --- | --- | --- |
|  | CTL | LIN | NIT | LIN+NIT |  | linseed | nitrate | lin*nit |  | CTL | TEA | NIT | TEA+NIT |  | saponin | nitrate | sap*nit |
| Entodinium | 59.84 | 49.66 | 46.52 | 33.94 | 0.036 | 0.67 | 0.05 | 0.10 |  | 46.21 | 51.05 | 47.55 | 50.55 | 0.040 | 0.68 | 0.90 | 0.88 |
| Polyplastron | 11.57 | 8.50 | 17.51 | 10.43 | 0.021 | 0.75 | 0.26 | 0.27 |  | 7.94 | 9.73 | 10.39 | 7.19 | 0.016 | 0.62 | 0.92 | 0.63 |
| Unclassified Trichostomatia | 9.93 | 11.59 | 15.26 | 16.70 | 0.023 | 0.97 | 0.36 | 0.75 |  | 22.28 | 9.34 | 14.59 | 14.16 | 0.032 | 0.23 | 0.72 | 0.47 |
| Isotricha | 1.52 | 3.92 | 6.00 | 4.34 | 0.009 | 0.15 | 0.06 | 0.68 |  | 5.43 | 5.27 | 6.72 | 6.43 | 0.008 | 0.56 | 0.65 | 0.76 |
| Dasytricha | 1.31 | 0.80 | 1.86 | 4.01 | 0.006 | 0.65 | 0.10 | 0.76 |  | 0.90 | 1.92 | 1.67 | 1.77 | 0.004 | 0.19 | 0.34 | 0.13 |
| Unclassified | 0.17 | 0.17 | 0.06 | 0.19 | 0.000 | 0.08 | 0.15 | 0.06 |  | 0.14 | 0.18 | 0.17 | 0.14 | 0.000 | 0.85 | 0.82 | 0.57 |
| Ophryoscolex | 0.16 | 0.00 | 0.46 | 1.33 | 0.002 | 0.06 | 0.00 | 0.67 |  | 3.26 | 3.08 | 0.41 | 1.47 | 0.010 | 0.64 | 0.52 | 0.78 |
| Unclassified Ciliophora | 0.04 | 0.09 | 0.04 | 0.06 | 0.000 | 0.59 | 0.50 | 0.27 |  | 0.03 | 0.07 | 0.06 | 0.05 | 0.000 | 0.09 | 0.81 | 0.09 |
| Unclassified SAR | 0.04 | 0.02 | 0.01 | 0.02 | 0.000 | 0.09 | 0.38 | 0.66 |  | 0.04 | 0.04 | 0.01 | 0.02 | 0.000 | 0.15 | 0.00 | 0.57 |
| Trichostomatia | 0.04 | 0.28 | 0.02 | 0.02 | 0.000 | 0.04 | 0.01 | 0.06 |  | 0.10 | 0.08 | 0.24 | 0.03 | 0.000 | 0.41 | 0.94 | 0.34 |
| Pseudoplatyophyra | 0.00 | 0.05 | 0.02 | 0.00 | 0.000 | 0.00 | 0.18 | 0.12 |  | - | 0.03 | 0.00 | 0.00 | 0.000 | 0.21 | 0.50 | 0.17 |

<sup>a</sup>In Study 1, cows were fed a control (CTL) diet and CTL diet supplemented with linseed (LIN), nitrate (NIT) and linseed plus nitrate (LIN+NIT). In Study 1, tested effects were linseed (CTL and NIT versus LIN and LIN+NIT) and nitrate (CTL and LIN versus NIT and LIN+NIT) and their interaction lin\*nit
<sup>b</sup>In Study 2, cows were fed a control (CTL) diet and control diet supplemented with tea saponin (TEA), nitrate (NIT) and tea saponin plus nitrate (TEA+NIT). In Study 2, tested effects were saponin (CTL and NIT versus TEA and TEA+NIT) and nitrate (CTL and TEA versus NIT and TEA+NIT) and their interaction sap\*nit.

### Tea saponins had only minor effects on rumen microbial population

Adding tea saponin to diets only affected the low-abundant order of unclassified *Deltaproteobacteria* (Table 2). No changes in diversity indices were noticed (Table S1). NMDS (Figure 3) and PERMANOVA analysis did not reveal significant changes in bacterial community, though *Lachnospiraceae* were highly abundant in cows supplemented with saponin (Figure 2). Similarly, concentration and taxonomic composition of the archaeal community were not influenced by tea saponin (Figure 3, Table 3), and neither was protozoa community structure (Figure 3, Table 4).

**Figure 3.**
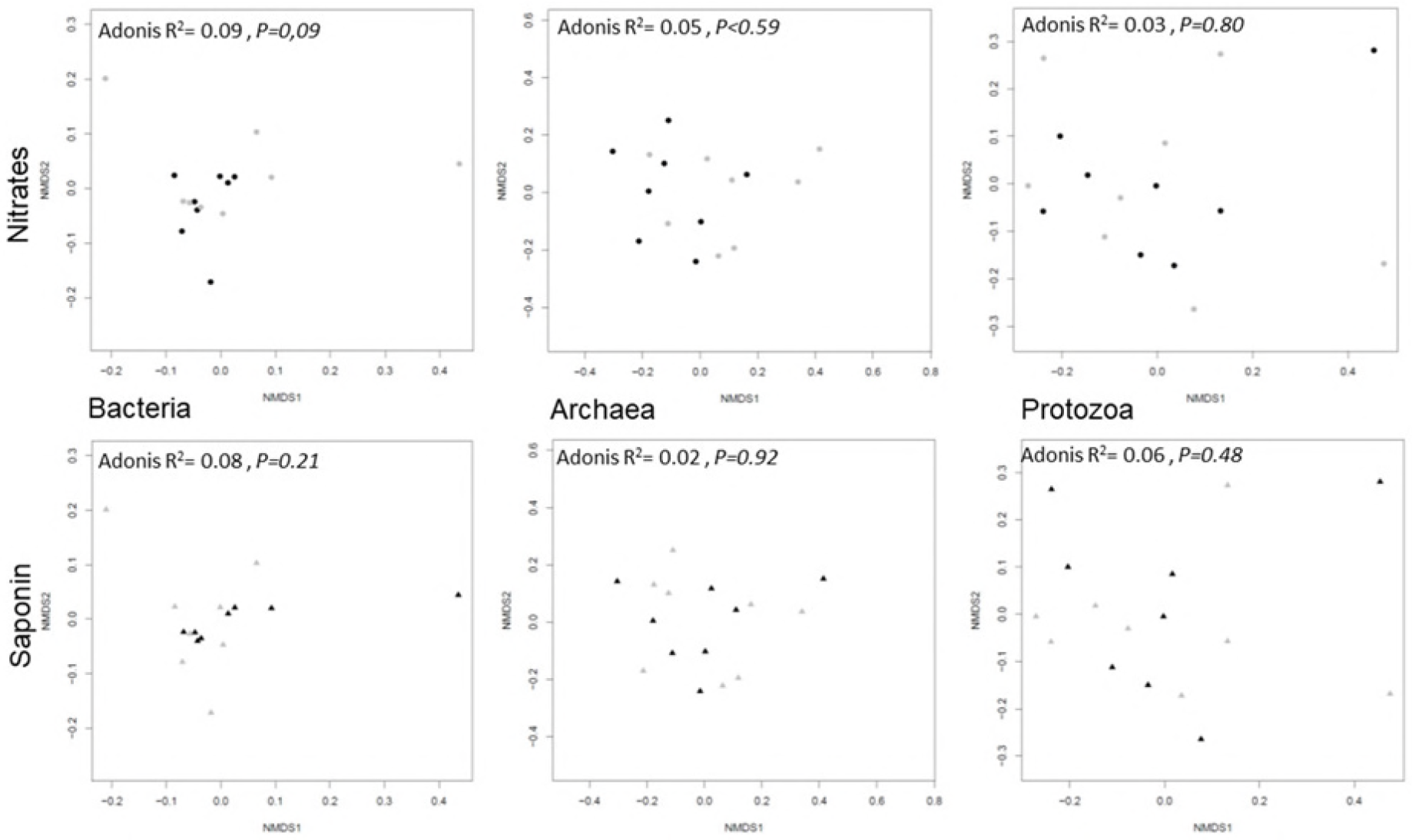
Structure and composition of bacterial, archaeal and protozoal communities in Study 2, related to nitrate or saponin treatments (black symbols) and respective controls (grey symbols), were examined by multivariate analysis. Non-metric multidimensional scaling plots derived from Bray Curtis dissimilarities between cows. Each symbol is representative of a single cow. Samples are plotted along the first two component axes. Microbial composition was compared using Adonis.

### Nitrate remodels bacterial and archaeal communities

In order to have an integrated discussion on the effects of nitrate on microbes from both studies, we needed to compare like to like. This is why we compared microbial communities of cows fed CTL-diet in each study. Bacterial communities of these cows were similar (Adonis R^2^ = 0.16 and *P* = 0.26). A small numerical difference was noted in the Bacteroidales:Clostridiales ratios, which were 1.02 and 0.81 in Study 1 and Study 2 respectively. As well as for bacteria, methanogenic communities in animals fed CTL diets were similar between studies (Adonis R^2^=0.028 and *P*=0.942). Regarding protozoa, some differences were revealed by NMDS and PERMANOVA analyses between the two control groups. NMDS graphs (Figure S2) showed only a small overlap between protozoa population fed CTL in each study, which was confirmed by an Adonis test (*P*<0.1). Also, *Entodinium-related* sequences accounted for 60% of total classified sequences in Study 1, whereas they represented 46% of sequences in Study 2 (Table 4). Though this difference was not statistically significant, it was accompanied by significantly higher numbers of *Trichostomatia* and *Isotricha* related sequences in Study 1 compared to Study 2 (Table 2).

Feeding nitrate, in both studies, increased *Coriobacteriales* and *Burkholderiales* abundance and decreased (Study 2) or tended to decrease (Study 1) abundance of *Gastranaerophilales* (Table 2). In addition, in Study 2, nitrate supplementation increased the relative abundance of *Bacteroidales* (Table 2). Diversity indices were not influenced by dietary treatment (Table S1) in any study. Non-metric multidimensional scale (NMDS) analysis (Figure 1) revealed that while in Study 1, nitrate supplementation was the major driver of phylogenetic dissimilarity among bacterial communities (Adonis R^2^=0.11, *P*<0.01), in Study 2 nitrate only moderately affected community structure (Adonis R^2^=0.09, *P*=0.09). Indicator species analysis revealed that 10 OTUs in Study 1 and 21 in Study 2 were differentially abundant between cows fed and not fed nitrate (P<0.05 and indicator value >0.7; Table S4 and Table S5). *Lachnospiraceae* and *Sutterellaceae* characterised nitrate-supplemented diets (Figure 2) in Study 1 and *Coriobacteriaceae* and uncultured *Mollicutes* family were identified as indicator OTUs for nitrate-supplemented diets in Study 2. More interestingly, in both studies *Ruminococcaceae*−related OTUs characterised the bacterial community of cows not fed nitrate (Figure 2).

In both studies, feeding nitrate had no effect on methanogen concentration in the rumen *(mcrA* copy numbers), but reduced methanogen activity *(mcrA* expression levels) (Table 1). When cows were fed nitrate, Shannon and Simpson diversity indices decreased or tended to decrease (Table S1), although the overall taxonomic composition was not affected (Table 3). NMDS and PERMANOVA analyses showed that feeding nitrates deeply modified archaeal community structure in Study 1, but had no effect on community structure in Study 2.

In Study 1, *Entodinium* relative abundance tended to decrease and *Isotricha* tended to increase in animals receiving nitrate-supplemented diets (Table 4). Diversity indices remained similar between diets and contrasts (Table S1). However, there was some evidence (Adonis R^2^=0.12, *P*=0.05) that nitrate modulated rumen protozoa population in cows (Figure 1). In contrast, in Study 2, nitrate had no effect on protozoa community in the rumen of non-lactating dairy cows (Figure 3).

### Correlation patterns of microbial population

We analysed the correlation between bacterial families and genera of methanogens and protozoa (Figure 4 and Figure 5). Values for methane production (g/day), yield (g/kg DMI), hydrogen production (only for Study 1) and (acetate+butyrate)/propionate ratio from the datasets of Guyader et al (8, 13) were also included in the analysis. Only significant correlations are discussed.

**Figure 4.**
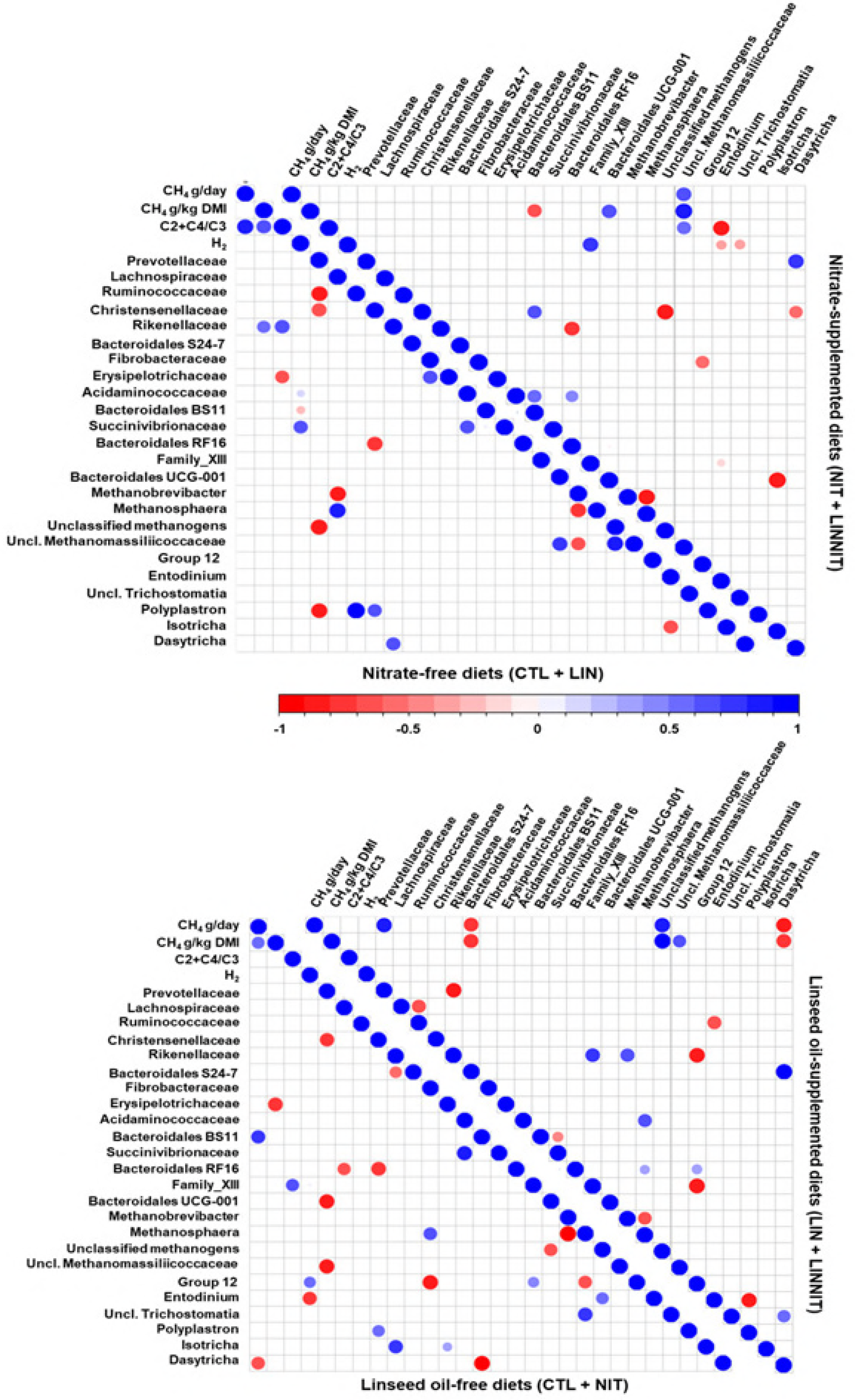
Spearman’s rank correlation matrix of the dominant ruminal bacterial families, genera of archaea and protozoa and fermentation parameters in Study 1. Illustrated correlation patterns are for nitrate and linseed supplementations. Listed microbial populations were detected in at least 50% of the rumen samples analysed and represent at least 1% of the bacterial, archaeal, protozoa, or fungal communities. Strong correlations are indicated by large circles, whereas weak correlations are indicated by small circles. The colours of the scale bar denote the nature of the correlation with 1 indicating perfect positive correlation (dark blue) and -1 indicating perfect negative correlation (dark red) between two microbial populations.

**Figure 5.**
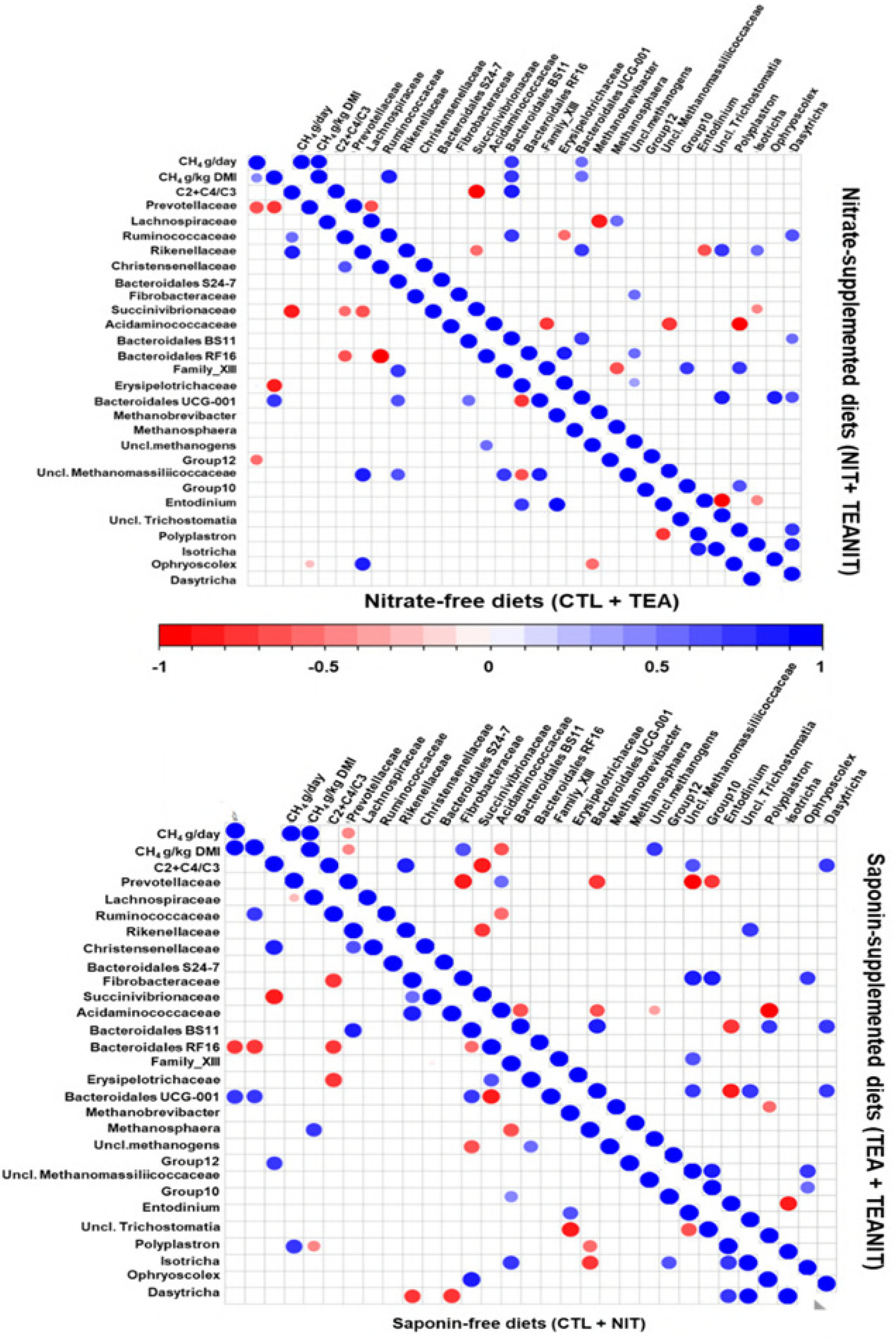
Spearman’s rank correlation matrix of the dominant ruminal bacterial families, genera of archaea and protozoa and fermentation patterns in Study 2. Illustrated correlation patterns are for nitrate and tea saponin supplementations. Listed microbial populations were detected in at least 50% of the rumen samples analysed and represent at least 1% of the bacterial, archaeal, protozoa, or fungal communities. Strong correlations are indicated by large circles, whereas weak correlations are indicated by small circles. The colours of the scale bar denote the nature of the correlation with 1 indicating perfect positive correlation (dark blue) and -1 indicating perfect negative correlation (dark red) between two microbial populations.

In Study 1 (Figure 4), methane production (g/day) and yield (g/kg DMI) were positively correlated (R^2^= 0.83, and R^2^= 0.69 respectively) with the (acetate+butyrate)/propionate ratio when cows were not fed nitrate; in these animals, methane yield correlated positively with *Rikenellaceae* (R^2^= 0.56). In the absence of nitrate, *Methanobrevibacter* negatively correlated with unclassified *Methanomassiliicoccaceae* (R^2^=-0.64) and *Ruminococcaceae* correlated positively with members of the protozoal *Polyplastron* genus (R^2^=0.97). When diets were supplemented with nitrate, methane production and yield as well as (acetate+butyrate)/propionate ratio were strongly correlated with a group of unclassified *Methanomassiliicoccaceae* (R^2^= 0.69, R^2^= 0.85 and R^2^=0.59 respectively). In addition, when diets were nitrate-supplemented, a positive correlation was established between *Prevotellaceae* and *Dasytricha* (R^2^= 0.70 and R^2^= 0.73). There was a strong negative correlation between *Methanobrevibacter* and *Methanosphaera* independently of nitrate supplementation (R^2^= -0.76 in cows not fed and R^2^= -0.83 cows fed nitrate).

Methane production and yield when linseed was fed to cows correlated negatively with *Bacteroidales* S 24.7 group (R^2^= -0.76) and *Dasytricha* (R^2^= -0.83) populations and positively with an unclassified archaeal taxon (R^2^= 0.83) and unclassified *Methanomassiliicoccaceae* (R^2^= 0.61). Independently of linseed supplementation, a negative correlation between *Methanobrevibacter* with *Methanosphaera* was observed (R^2^= -0.64 and R^2^= -0.95).

In Study 2 (Figure 5), when diets were not supplemented with nitrate, methane production correlated negatively with *Prevotellaceae* (R^2^= -0.69) and methanogen Group 12 (R^2^= -0.50). When diet was supplemented with tea saponin, *Prevotellaceae* correlated negatively with methane production (R^2^= -0.47) and yield (R^2^= -0.43), but also with *Fibrobacteraceae* (R^2^= -0.88), *Bacteroidales* (R^2^= -0.76) and two families of *Methanomassiliicoccaceae* (R^2^= -0.90 for unclassified *Methanomassiliicoccaceae* and R^2^= -0.70 for Group 10).

## Discussion

Guyader et al., (8) showed that combining dietary strategies acting theoretically on hydrogen production (lipids) and consumption (nitrate) can have an additive effect on methane reduction. In a second study, they confirmed the antimethanogenic potential of nitrate supplementation, but observed no effect of tea saponin on methane production (13). These studies were conducted simultaneously; cows were selected at random from the same experimental herd and were randomly allocated to a study. Given the consistency of results on methane production and fermentation patterns reported in the two articles of Guyader et al (8, 13), we decided to analyse the rumen microbiota from both studies at the same time (from DNA extraction up to statistical tests). Although linseed and nitrate have a medium to high potential methane mitigating effect (the effect of saponins being less reproductive) (19), microbial data is scarce and inconsistent between studies. This could be explained by different methodologies for rumen sample collection, conservation and nucleic acid extraction, but also on how data was obtained and analysed (20, 21). Thus, secondly, we compiled the microbial data in order to get an insight into the mode of action of nitrate, linseed and saponin on the rumen microbial ecosystem.

To this aim, we first checked that microbiota of the two groups of cows was comparable; hence, we performed a detailed analysis of microbial community structure and composition in rumen contents sampled during the period when CTL diet was fed to each animal. No major differences in bacterial communities were observed, except a non-significant shift in Bacteroidales:Clostridiales ratio, which is known to vary widely across individual animals (22). However, we observed numerical differences in the relative abundance of *Entodinium* (60% in Study 1 *vs*. 46% in Study 2), which is consistent with enumeration results reported previously (5.71 and 5.38 log_10_ cells / mL in Study 1 and Study 2 respectively (8, 13) showing more abundant ciliate populations in cows from Study 1.

In Study 1, dietary supplementation with linseed increased the relative abundance of *Selenomonadales*. This is in accordance with our previous work exploring the effects of linseed plus nitrate on rumen microbiota (23) in bulls, where we reported increased numbers in sequences affiliated with three *Selenomonas* genera and one unclassified *Selenomonadales* genus. As these microbes are potential nitrate reducers (24), we hypothesised that their growth was supported by the higher nitrate availability, but the present study suggests that it is rather a linseed effect. Oleic acid (representing on average 20% of linseed oil fatty acids) stimulated the growth of *Selenomonas ruminantium* in pure cultures (25). However, for *in vivo* studies, results are contrasting: *Selenomonas* was among the genera explaining differences in bacterial community structure between lambs fed linseed diet and those fed control diet (26); but there was no change in *Selenomonas* abundance when cows were fed sunflower oil (30% oleic acid) (22). Members of the *Selenomonadales* order are also known to reduce succinate to propionate, which is in agreement with higher molar proportion of propionate in the rumen of cows fed linseed (8). Linseed supplementation also increased abundance of uncultured *Bacteroidetes* and the *Bacteroidales* S27-7 family was negatively correlated with methane production and yield. On the other hand, linseed diets were characterised by decreased abundance of *Ruminococcaceae*, which is in agreement with previous findings that fatty acids are toxic to these cellulolytic microbes (22, 25, 27). We observed no effect on rumen protozoa numbers (8) and diversity, though *Dasytricha* correlated negatively with methane emissions and positively with the *Bacteroidales* S27-7 family. Linseed oil supplementation also had no effect on the abundance or diversity of the rumen methanogenic community. In accordance with previous results (9, 23), the antimethanogenic potential of linseed oil fatty acids was not related to archaeal numbers in the rumen, but rather to a lower metabolic activity of these microbes, which could be explained by lower availability of hydrogen.

Adding tea saponins to the diet had no effect either on microbial numbers or on diversity. This is consistent with the lack of changes in methane production or VFA profiles reported by Guyader et al (13). The efficacy of saponins in suppressing methane production varies considerably depending on the chemical structure, source, dose, and diet (28). Saponins have been reported to inhibit rumen protozoa (5) and thus limit hydrogen production in the rumen. However, in our previous work (13) and the study of Ramírez-Restrepo (29), adding tea saponins to ruminants’ diets had an opposite effect on protozoal numbers. Saponins break down the membrane of protozoa by interacting with its sterols. However, rumen microbes can degrade the sugar moiety of saponins rendering them inactive. Recently it was proposed to improve the antiprotozoal effect of saponins by changing their chemical structure and thus protecting them from microbial degradation (12).

In Study 1, multivariate analysis revealed that nitrate supplementation altered bacterial and archaeal communities. However, in Study 2, NMDS and PERMANOVA results were less conclusive, though reductions of methane emissions and changes in fermentation parameters were comparable between experiments. Nevertheless, both studies pinpointed a limited number of taxa associated with decreased methane emissions in nitrate-fed cows. Nitrate supplementation increased the abundance of *Coriobacteriales* and *Burkholderiales* orders, which contain taxa with known nitrate-reducing activity (30-32). This is in accordance with the numerically higher nitrite concentrations measured by Guyader et al (8, 13) in nitrate-fed cows. Also, cows not fed nitrate presented an enhanced cellulolytic community, which is in accordance with our previous results showing a toxic effect on *Ruminococcaceae* in animals fed linseed plus nitrate diets (23). *Ruminococcus flavefaciens* and *Ruminococcus albus* populations decreased in the rumen of goats when nitrate was added to the diet (24). An *in vitro* study (33) showed that the growth of these two cellulolytic bacteria was inhibited by nitrite at a level of 3 mmol/L, but measured nitrite levels in our studies rarely exceeded 0.08 mmol/L (8, 13). Lower concentrations could still be toxic, as another study showed that the specific growth rate of *R. flavefaciens*, but not *R. albus*, was decreased by less than 0.03 mmol/L of nitrate (34). Marais et al.(34) also argued that nitrite inhibits electron transport systems (*R. flavefaciens*), so bacteria not possessing an electron transport system (*R. albus*) are less affected. *R. flavefaciens* and *R. albus* are the only cultured *Ruminococcus* species able to degrade cellulose (35), making them an important part of a functional rumen ecosystem. *In vitro, R. albus* produces acetate, hydrogen and carbon dioxide and its metabolic activity is stimulated by the presence of methanogens (36). Reducing *Ruminococcaceae* numbers by nitrate supplementation would thus decrease the amount of hydrogen produced, which could indirectly reduce methane production. This conclusion is also supported by the decreased expression levels of the methanogen *mcrA* gene, which has been shown to correlate with methane emissions (23, 37, 38). However, *Ruminococcaceae* are an important group of bacteria inhabiting the rumen and are able to degrade plant cell wall polysaccharides into metabolisable energy. This would imply that inhibition of the rumen fibrolytic community is likely to decrease fibre degradation. In the present studies, nitrate supplementation did not affect total tract digestibility (8, 13), but linseed tended to reduce fibre digestibility (8).

We also observed a strong positive correlation between unclassified *Methanomassiliicoccaceae* and methane production when cows were fed nitrate-supplemented diets. Veneman et al (9) also reported an increase in the abundance of *Methanomassiliicoccaceae*−related methanogens in the rumen of nitrate-fed animals. *Methanomassiliicoccaceae* are obligate hydrogen-dependent methylotrophic methanogens (39), whereas most of the other rumen methanogens perform methanogenesis from hydrogen and carbon dioxide. They are part of a unique methanogen order with a characteristic set of genes involved in the methanogenesis pathway (39). It is likely that their particular physiology confers on them a competitive advantage when the activity of other methanogens is affected in a nitrate/nitrite-enriched environment.

We conducted this study to understand how the rumen microbial ecosystem responds to dietary methane mitigation by linseed, saponin and nitrate supplementation alone or in combination. We hypothesised that adding linseed or saponins to the diet reduces hydrogen production by a toxic effect on rumen protozoa and by replacing dietary carbohydrates by non-fermentable fatty acids; additionally, we were expecting that nitrate supplementation would redirect hydrogen consumption towards nitrate reduction, rather than methanogenesis. Changes in the rumen microbial ecosystem were monitored using archaea, bacteria and eukaryotes specific primers targeting either 16S or 18S rRNA. Our sequencing strategy allowed us to accurately draw the parallel between changes in methane emissions and microbiota structure. Our study showed that linseed oil decreases methane emissions by reducing the number of hydrogen producers (cellulolytic *Ruminococcaceae)* and by stimulating propionate producers *(Selenomonas)*, thus diverting hydrogen from methanogenesis. Nitrate supplementation favoured the development of nitrate-reducing bacteria *(Coriobacteriales* and *Burkholderiales)* and had a negative effect on cellulolytic *Ruminococcaceae*; as a consequence, nitrate supplementation also significantly affected methanogen community structure and activity. We did not show any shifts in rumen microbiota structure and activity due to dietary supplementation with tea saponins.

In a secondary aim of our work, we capitalised on data available from two independent studies expecting to draw relevant conclusions. It is common that studies exploring microbial mechanisms of the same methane abatement strategy come to dissimilar conclusions. Authors generally argue that these differences are due to differences in diet, animal species, physiologic stage, and different sample processing or bioinformatics pipelines. In the present work, we minimised the impact of study design on data interpretation, despite some inconsistent results were observed for nitrate-supplemented diets from Study 1 and Study 2. Nitrate reduced methanogen activity and stimulated nitrate-reducing bacterial populations in both studies. Similarly, *Ruminococcaceae*−related OTUs characterised nitrate-free diets in both studies. In contrast, multivariate analysis showed that nitrate altered bacterial and archaeal communities in Study 1, whereas only a moderate effect on bacteria was observed in Study 2. In both experiments each experimental period lasted 5 weeks. It is possible that microbiota shifted as a result of imposed dietary treatments and did not completely migrate back to the initial state. In a massive contents exchange trial, Weimer et al., (40) found that cows reconstructed almost completely their microbiota in 3 weeks, with a complete return to its original host-specific state in 9 weeks. However, pH and VFA profiles returned to the original values much more quickly, within one day. We could argue that changes induced by nitrate supplementation were at the level of microbe function rather than species composition. This is supported by the fact that reductions in methane emissions and shifts in VFA profiles were comparable between studies. A metatranscriptomic approach will be more fruitful to further explore microbial mechanisms of methane mitigation using linseed and/or nitrate.

## Materials and methods

The experiments were conducted at the animal facilities of INRA Herbipôle Unit (Saint-Genès Champanelle, France). All procedures involving animals were conducted in accordance with the French Ministry of Agriculture guidelines for animal research and all applicable European guidelines and regulations on animal experimentation. The experiments were approved by the Auvergne Regional Ethics Committee for Animal Experimentation, approval number CE50-12.

### Animals, experimental design and feeding management

Animals and experimental design are those described in Guyader et al., 2015 a and b (8, 13). Briefly, eight non-lactating Holstein cows were separated into two groups conducted in parallel according to a 2 × 2 factorial design. Within each study, four cows were randomly assigned to four dietary treatments during 5-week experimental periods. In Study 1, diets were on a DM basis: 1) control diet (CTL, 50% natural grassland hay and 50% concentrate), 2) control diet with 4% linseed oil (LIN; 2.6% added fat), 3) control diet with 3% calcium nitrate (NIT; 2.3% nitrate), and 4) control diet with 4% linseed oil plus 3% calcium nitrate (LIN+NIT; 2.6% added fat plus 2.3% nitrate)(8). In Study 2, diets were on a DM basis: 1) control diet (CTL, 50% natural grassland hay and 50% concentrate), 2) control diet with 0.77% tea saponin (TEA; 0.5% saponin), 3) control diet with 3% calcium nitrate (NIT; 2.3% nitrate), and 4) control diet with 0.77% tea saponin plus 3% calcium nitrate (TEA+NIT; 0.5% saponin plus 2.3% nitrate)(13). The chemical compositions of the diets CTL and NIT were similar between the two studies.

### Rumen content sampling for microbial analysis

At the end of each experimental period, whole rumen content samples (200 g) were taken, through cannula, from multiple sites within the rumen. Sampling was done 3 h after the morning feeding when methane emissions differences between diets measured in the same animal were maximal (41). A part of each sample (~30 g) was mixed with 30 mL ice cold PBS pH 6.8 and homogenised using a Polytron grinding mill (Kinematica GmbH, Steinhofhalde, Switzerland) for three cycles of 1 min with intervals of 1 min on ice. Then, approximately 0.5 g was transferred to a 2.5 mL Eppendorf tube and mixed with 1 mL of RNA*later*^®^ Stabilization Solution (Applied Biosystems, Austin, TX, USA). Tubes were immediately stored at -80°C until further processing.

### Total nucleic acid extraction and cDNA synthesis

Total nucleic acids (DNA and RNA) were co-extracted from all samples by bead-beating and phenol-chloroform extraction followed by saline-alcohol precipitation (42). The yield and purity of extracted DNA and RNA were assessed using a Nanodrop Lite Spectrophotometer (Thermo Fisher Scientific, Wilmington, USA); RNA integrity was estimated with an Agilent RNA 6000 Nano Kit on an Agilent 2100 bioanalyser (Agilent Technologies, Santa Clara, CA, USA) according to the manufacturer’s instructions. Following extraction and quality assessment, RNA was reverse-transcribed using the Reverse Transcriptase Kit with random primers (Promega, Madison, USA) according to the manufacturer’s instructions, on a T-100 thermocycler (BioRad, Hercules, USA).

### Quantification and gene expression of microbial communities

Samples from each cow from the two sampling days of each experimental period were pooled by mixing an equal quantity of DNA or equal volumes of cDNA. Quantification of gene targets was performed on microbial DNA and cDNA by quantitative PCR (qPCR) using a Step One Plus apparatus (Applied Biosystems, Villebon sur Yvette, France). Reactions were run in triplicate in 96-well plates, using 15.5 μL of 1X Takara SYBR Premix Ex Taq (Lonza, France), 0.25 μmoles of each forward and reverse primer and 20 ng of DNA or 2 μL of cDNA in a final volume of 20 μL. Primer description, average amplification efficiency, slope and R^2^ of qPCR are described in Table S4 as required by MIQE guidelines for PCR (43). Negative controls without templates were run in each assay to assess overall specificity.

Abundances of total bacteria (based on 16S rDNA copies) and methanogens (based on *mcrA* DNA copies) were assessed using absolute quantification as previously described (38). Level of expression of the functional *mcrA* gene (based on *mcrA* cDNA copies) was assessed using the 2^-ΔCt^ method (44) with 16S rDNA copies as internal reference.

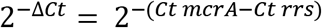

Technical triplicates were averaged while checking overlaying of amplification plots at threshold cycle (C_t_) value. Absolute quantification of total bacteria and methanogenic archaea was expressed respectively as log_10_ 16S rDNA or *mcrA* copies/ng extracted DNA.

### Sequencing strategy and data analysis

Approximately 3 μg of extracted DNA were sent to the Roy J. Carver Biotechnology Center (Urbana, IL61801, USA) for fluidigm amplification and sequencing of bacterial and archaeal 16S rDNA, and eukaryotic 18S rDNA for protozoa (Table S4). The libraries were sequenced on a 250 paired-ends MiSeq run and generated 8 249 698 raw reads for bacterial 16S rDNA, 1 778 521 for archaeal 16S rDNA and 2 245 531 for eukaryotic 18S rDNA (Table S5). Data was analysed on a home-made Galaxy-based graphic user interface for QIIME (45) and IM Tornado (46) (Table S1). All pipelines included a quality control step, removing sequences with phred score <33 and trimming based on expected amplicon length, merging paired reads, chimera search and removal and OTU picking (Table S2). Merging paired end 16S rDNA reads was performed by mothur’s (47) make.contigs command before input in the QIIME pipeline. Taxonomic classification for bacteria and protozoa was based on SILVA v123 database (48) and for archaea on RIM-DB (49).

### Statistical analysis

Results from qPCR quantification, relative abundance (after square root transformation) of microbes at different taxonomic levels and diversity indices were analysed by ANOVA in R(version 3.4.0). The statistical model included the random effect of cow (n = 4) and fixed effects of period (n = 4) and contrasts for nitrate (CTL and LIN *versus* NIT and LIN+NIT in Study 1; CTL and TEA *versus* NIT and TEA+NIT in Study 2), linseed (CTL and NIT *versus* LIN and LIN+NIT in Study 1), tea saponin (CTL and NIT *versus* TEA and TEA+NIT in Study 2) and the interactions linseed × nitrate or saponin × nitrate. Significance was considered at *P*≤0.05. Trends were discussed at 0.05<*P*≤0.1. Least square means are reported throughout.

OTUs with fewer than 3 sequences were withdrawn from further analysis. OTU tables were imported in R and rarefied to minimise the variations created by different sample depth subsampling. Further analysis was performed using the vegan R package (50). Alpha diversity values for all microbial communities were obtained using various diversity indices (Shannon and Simpson diversity indices, richness and evenness) and analysed by ANOVA for the effect of contrasts and the interactions described above. A nonmetric multidimensional scaling was used to ordinate microbial libraries (n=4*4 per study and per microbial group). We used the betadisper function to check the homogeneity of group dispersions before performing a PERMANOVA analysis via the Adonis function of vegan. The multipatt function from R package indicspecies (51) was used to find indicator OTUs using a 5% significance level for selecting indicators in cows fed linseed, tea saponin and nitrate. The species-site group association parameter was “IndVal.g”.

Correlation analysis between microbial populations and some fermentation parameters (methane, hydrogen, VFA ratio) were performed in R. Only microbial groups that represented more than 1% (average of all samples) of the total community within each of the three microbial groups (bacteria, archaea or protozoa) and that were detected in at least 50% of rumen samples were included in the analysis. Spearman’s rank correlations and P-values were calculated by above described contrasts and plotted using the packages hmisc (52) and corrplot (53).

## Acknowledgements

J. Guyader was the recipient of an INRA-Région Auvergne PhD scholarship. C. Saro acknowledges receipt of a postdoctoral fellowship from Fundación Alfonso Martín Escudero (Madrid, Spain). The authors also thank NEOVIA for financial support. The authors thank the INRA personnel of the Experimental Unit Herbipôle (L. Mouly, D. Roux, S. Rudel and V. Tate) for taking care of the animals, and of the UMR Herbivores (L. Genestoux and D. Graviou) for their help in sampling and performing laboratory analysis.

The authors thank David Marsh for checking and amending our English

We are grateful to Prof Richard Dewhurst for constructive criticism of the manuscript.

## Author Contributions

Conception or design of the work: MP, JG, MS, DM. Data collection: MP, JG, MS, CS, ARS. Data analysis and interpretation: MP, JG, MS, CS, ARS, CG, CM, DM, AB. Drafting the article: MP. Critical revision of the article: JG, MS, CS, ARS, CG, CM, DM, AB. Final approval of the version to be published: MP, JG, MS, CS, ARS, CG, CM, DM, AB.

## Additional Information

Raw sequence data is available in the Sequence Read Archive (SRA) under BioProject ID PRJNA415383.

**The authors declare no competing financial interests**.

